# MiRNAs profiling and degradome sequencing between the CMS-line N816S and its maintainer line Ning5m during anther development in pepper (*Capsicum annuum* L.)

**DOI:** 10.1101/2020.02.04.933473

**Authors:** Hongyuan Zhang, Shuping Huang, Jie Tan, Xia Chen, Min Zhang

**Author notes:** Corresponding author: Min Zhang.

## Abstract

Utilization of cytoplasmic male sterility (CMS) is significant for agriculture. MiRNAs are a class of endogenously non-coding small RNAs (21-24 nt) that play key roles in the regulation of various growth and developmental processes in plants. The knowledge miRNA-guided CMS regulation is rather limited in pepper. To better understand the miRNAs involvement and regulatory mechanism of CMS, miRNA libraries from anther of CMS-line N816S and its maintainer line Ning5m were generated by miRNAome sequencing in pepper. A total of 76 differentially expressed miRNAs were detected, of which 18 miRNAs were further confirmed by quantitative real-time PCR (qRT-PCR). In addition, miRNA targets were identified by degradome sequencing. The result showed that 1292 targets that were potentially cleaved by 321 miRNAs (250 conserved miRNAs and 71 novel miRNAs). Gene Ontology (GO) and KEGG pathway analysis indicated that 35 differentially expressed miRNAs might play roles in the regulation of CMS sterility, by cleaving 77 target transcripts, such as *MYBs, SPLs*, and *AFRs*, of which targeted by miR156, miR167, miRNA858 family. Nineteen miRNA-cleaved targets were selectively examined by qRT-PCR, and the results showed that there were mostly negative correlations between miRNAs and their targets on the expression level. These findings provide a valuable information to understand miRNAs mechanism during anther development and CMS occurrence in pepper.

## Introduction

MicroRNAs (miRNAs) are a class of endogenous non-coding small RNAs (sRNAs) with 21-24 nucleotides (nt) in animals and plants(Zhou et al., 2020). In plants, mature miRNA sequences were generated from primary miRNA transcripts (pri-miRNA) by Dicer-like (DCL1)(Zhang et al., 2015). The mature miRNAs are incorporated into the RNA-induced silencing complex (RISC) to guiding the cleavage of specific complementary mRNAs(Schuck et al., 2013). MiRNAs are widely considered as negative-regulated gene expression at the post-transcription or translation process via degrading target mRNAs or repressing mRNA translation(Stepien et al., 2017). Previous studies have manifested that miRNAs involved in various plant growth and developmental processes, such as hormone homeostasis, flower development, embryogenesis and stress responses(Jha and Shankar, 2011;Pei et al., 2013).

Recently, a large number of studies have revealed miRNAs as important regulators of gene expressions to be involved in the developmental transition from vegetative growth to reproductive growth(Lin et al., 2013). MiR156 and miR172, two prime participators in flowering regulation, have been shown to regulate the floral transition in plant(Zhu and Helliwell, 2011;Yu et al., 2012;Diaz-Manzano et al., 2018). MiR156 could target SQUAMOSA Promoter Binding Protein-like (SPL) Transcription Factors (TF) and decreases with increasing life spans of the plant(Zheng et al., 2019). Overexpression of miR156 could delay flowering and prolong vegetative stage in many species, including Arabidopsis, rice and tomato(Zheng et al., 2019). The miR172, which targets APETALA2-like (*AP2*) transcription factors, has the opposite effect to the miR156 on the regulation of flowering time and increases with phase development and induces floral organogenesis and promote flowering(Tripathi et al., 2018). In addition, miR397 has been reported promoting panicle branching, increasing grain size, and resulting in improving yield in rice(Zhang et al., 2013). These studies showed that miRNAs play important roles in a number of developmental processes and pathways which regulate flower development related process.

Pepper (*Capsicum annuum* L.) is one of the most economically important worldwide vegetable crops(Barrajon-Catalan et al., 2020). Hybrid breeding has made a tremendous contribution to pepper yield, increasing seed production efficiency and protection of the varieties patent(Jifon et al., 2019). The utilization of male sterility in hybrid pepper is mainly based on three-line systems, which including cytoplasmic male sterile line (CMS), a maintainer line and restorer line(Bohra et al., 2016). In the CMS pepper, an ORF named *orf456*, was found at the 3′-end of the *coxII* gene, which is concluded that the *orf456* may represent a candidate gene, from mitochondrial genes, for determining the male-sterile phenotype of CMS(Kim et al., 2007). Furthermore, serval CMS-related sterile genes and fertility restorer genes also have been cloned from various plants(Bohra et al., 2016). In Arabidopsis, miR167 overexpression has been reported leading to male fertility defects(Ru et al., 2006), whereas miR159a overexpression results in decreased expression of *MYB33* and *MYB65*, leading to male sterility and delays flowering time(Anthony, 2005). With the development of miRNAome sequencing technology, differential expression patterns of miRNAs between the cytoplasmic male sterility (CMS) line and its maintainer line have been reported in many vegetable crops, such as Brassica juncea(Yang et al., 2013), Chinese cabbage(Wei et al., 2015) and Radish(Zhang et al., 2016b). A large number of miRNAs related to flowering and flower development have been identified and characterized in above species. Nevertheless, there are no reports on systematic identification and characterization of CMS-related miRNAs in pepper.

To explore the roles of miRNA in CMS, we identified the miRNAs via a high-throughput sequencing approach from pepper anthers at the early uninucleate stage of the sterile line N816S and its maintainer line Ning5m. Differential expression patterns of miRNAs were analyzed and compared between N816S and Ning5m. Targets were predicted by degradome sequencing. These results may provide insights into clarification of the molecular mechanisms underlying the regulation of miRNAs during pollen development.

## Materials and Methods

### Pepper materials

The CMS sterile line N816S and its maintainer line Ning5m were used in this study. The plants were grown in greenhouse of the vegetable institute of Wuhan academy of agricultural sciences under normal conditions. In general, anthers at the uni-nucleate stage were manually collected. Anthers were harvested from three individual plants of each cultivar, immediately frozen in liquid nitrogen, stored at −80°C, and then used for RNA isolation. The microspore development was judged by both the floret length as described by Parra- Vega et al (Parra-Vega et al., 2013).

### Small RNA library Construction, sequencing, and miRNA analysis

Total RNA was extracted using Trizol reagent (Invitrogen, CA, USA). RNA quantity was detected by Qubit Fluorometer (Invitrogen, CA, USA)), and RNA purity was assayed by NanoDrop spectrophotometer (BioRad, PA, USA) and Agilent2100 bioanalyzer (Agilent Technologies, CA, USA). According to the manufacturer’s instructions, sRNA libraries were constructed using the Small RNA Sample Prep Kit (Illumina, CA, US) and meanwhile the sRNA passed quality test(Yeri et al., 2018). Briefly, the small RNA (18-30 nt) were ligated to a 3′ adaptor and a 5′ adaptor sequentially and then converted to cDNA by RT-PCR. The purified cDNAs after reverse transcription reaction were sequenced by the Illumina Hiseq2000 (Illumina, CA, USA). After the Illumina sequencing, the raw sequences were obtained through a quality control process to generate high quality reads, and clean reads were directly used for further bioinformatics analysis with ACGT101-v4.2-miR (LC Sciences, TX, USA) to remove adapter sequences, junk reads, short reads, common RNA families (rRNA, tRNA, snRNA and snoRNA), repeats(Jeyaraj et al., 2019). Because of lacking known miRNA records of pepper in miRNA database (miRBase 21 released), unique sRNA sequences (18-25 nt) were mapped to specific species precursors in the miRBase 21 and the pepper reference sequence to identify conserved miRNAs by BLAST search. New miRNAs were identified by extracting flanking genome sequence of unique sRNAs using MIREAP (http://sourceforge.net/projects/mireap/) (Huang et al., 2010), followed by the prediction of secondary structures by Mfold program (http://unafold.rna.albany.edu/?q=mfold) (Reuter and Mathews, 2009).

### Degradome Library Construction, Data analysis and Target identification

The degradome library, a mixed samples’ library, was constructed according to the method described previously using sliced ends of polyadenylated transcripts(German et al., 2009). In brief, poly-A-containing mRNA was purified from total RNA mixture of N816S and Ning5m, then ligated to 5’ RNA adaptor containing a *Mme*I recognition site. Subsequently, RT-PCR was performed to first-strand cDNA, followed by digestion with *Mme*I, and then ligated to 3’ adaptor. Finally, ligation product was amplified, purified and subjected to Illumina sequencing.

Raw reads were performed to remove adaptor sequence and low-quality reads resulting in clean reads. The high-quality specific sequences of were collected for subsequent degradome tags analysis. To identified potentially sliced targets of miRNAs, degradome sequence analysis were processed using the Cleveland 3.0 software package and the ACGT301-DGE program (LC Sciences, TX, USA)(Gong et al., 2015). The tags, which mapped to sense cDNA, were used to predict cleavage sites. Height of the degradome peak at each occupied transcript position was placed into five possible categories.

### Quantitative real-time PCR (qRT-PCR) validation

Total RNA was extracted from pepper anthers, and RNA-free DNase I (Fermentas, USA) was used to remove DNA contamination for 15 min at 37°C. Stem-loop qRT-PCR were carried out to validate differential expressional levels of miRNAs. The mRNA template for the miRNA target was reverse transcribed using the OligodT_20_ primer for qRT-PCR. All miRNA detection primers were designed and synthesized based on the mature miRNA sequences. For each miRNA, approximately 1 μg of total RNA was reverse-transcribed by reverse transcriptase using miRNA-specific stem-loop primers and a Fermentas Revert Aid First Strand cDNA Synthesis Kit (Fermentas, USA). Relative expression analysis of the miRNA and its target were performed using the ABI Step One Plus™ Real Time PCR System (Applied Biosystems, USA) and SYBR Green Master Mix (Roche, Germany). All reactions were run with three individual biological replicates, and *18S rRNA* was used as the internal control gene refer to Hwang et al(Hwang et al., 2013). The relative expression of miRNA and mRNA were used quantified the 2^−ΔΔCt^ method to calculate the fold change between N816S and Ning5m(Asha et al., 2016). The primers used are listed in Table S1.

## Results

### Overview of Small RNAs Libraries Sequencing Date

To determine the involvement regulatory roles of miRNAs in the fertility of sterile and maintainer lines during anther development in pepper. Six small RNA libraries, including three biological replicates from N816S (N816S_1, N816S_2, N816S_3) and Ning5m (Ning5m_1, Ning5m_2, Ning5m_3), were constructed for deep sequencing. An average of 14,567,767 and 14,274,100 raw reads were obtained from N816S and Ning5m anthers, respectively, which after length, Junk reads, Rfam, Repeat, mRNA, rRNA, tRNA, snoRNA and snRNA reads filtering, an average of 10,780,898 (72.21%) valid reads representing 6,882,786 (84.68%) unique sequences and 10,442,445 (74.25%) valid reads representing 6,666,094 (86.4%) unique reads, respectively (Table 1).The proportion of the valid reads in the corresponding raw reads was more than 70%, which suggested that the quality of the sequencing data was high (Table1). The length of total sRNAs and ranged from 18 to 25 nt, and in both the N816S and Ning5m libraries, the 24 nt category was most abundant (average of 55.83% and 59.92% in N816S and Ning5m libraries, respectively) (Figure 1A-B). The length of unique sRNAs, in both the N816S and Ning5m libraries, the 24 nt category was most abundant (average of 59.15% and 62.90% in N816S and Ning5m libraries, respectively) (Figure 2C-D). This is consistent with the typical lengths of plant sRNAs reported in other studies(Asha et al., 2016;Hu et al., 2016).

**Table 1.** Overview of reads from raw data to cleaned sequences.

| Category | N816S_mean |  | Ning5m_mean |  |
| --- | --- | --- | --- | --- |
|  | Total sRNAs (%) | unique sRNAs | Total sRNAs (%) | unique sRNAs |
| Raw reads | 14567767 (100%) | 8074144 (100%) | 14274100 (100%) | 7568101 (100%) |
|  |  | 1003388 |  |  |
| 3ADT&Length filter | 1255570 (8.62%) | (12.43%) | 1012608 (7.09%) | 707822 (9.35%) |
| Junk reads | 74827 (0.51%) | 54461 (0.67%) | 74748 (0.52%) | 54565 (0.72%) |
| Rfam | 328527 (2.26%) | 10516 (0.13%) | 382951 (2.68%) | 11080 (0.15%) |
| mRNA | 2437774 (16.73%) | 132754 (1.64%) | 2728370 (19.11%) | 138920 (1.84%) |
| Repeats | 8437 (0.06%) | 306 (0.00%) | 8898 (0.06%) | 293 (0.00%) |
|  | 10780898 | 6882786 | 10442445 | 6666094 |
| Valid reads | (74.01%) | (85.24%) | (73.16%) | (88.08%) |
| rRNA | 254459 (1.75%) | 8362 (0.10%) | 323537 (2.27%) | 9018 (0.12%) |
| tRNA | 47680 (0.33%) | 833 (0.01%) | 32260 (0.23%) | 786 (0.01%) |
| snoRNA | 2925 (0.02%) | 191 (0.00%) | 2198 (0.02%) | 165 (0.00%) |
| snRNA | 2671 (0.02%) | 182 (0.00%) | 1554 (0.01%) | 115 (0.00%) |
| other Rfam RNA | 20793 (0.14%) | 948 (0.01%) | 23402 (0.16%) | 997 (0.01%) |

**Figure 1.**
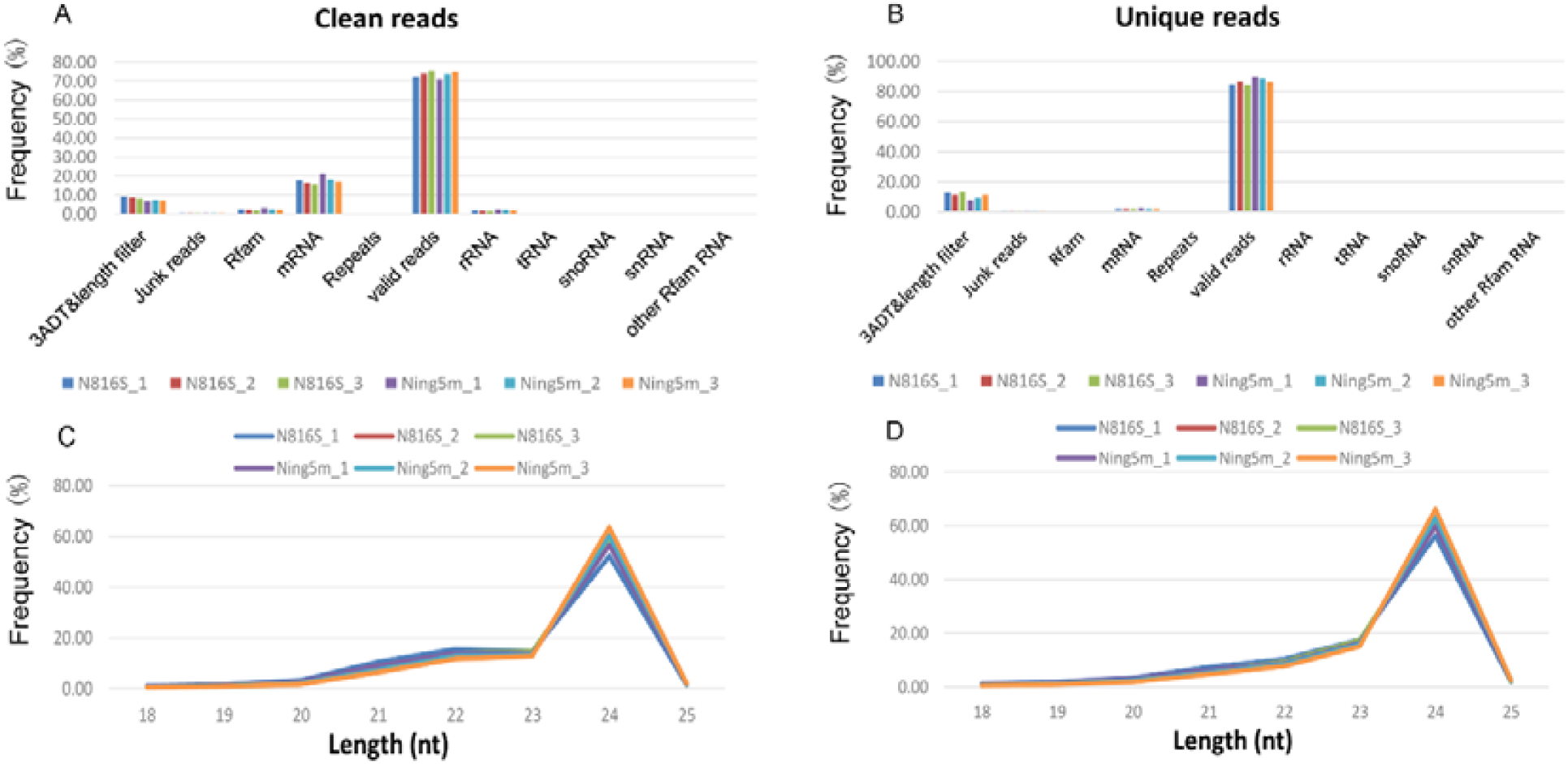
Frequency percent of Clean reads (A) and Unique reads (B)sequencing reads, and the length distribution of the clean reads (C) and unique reads (D) from anther of Capsicum annuum L.

**Figure 2.**
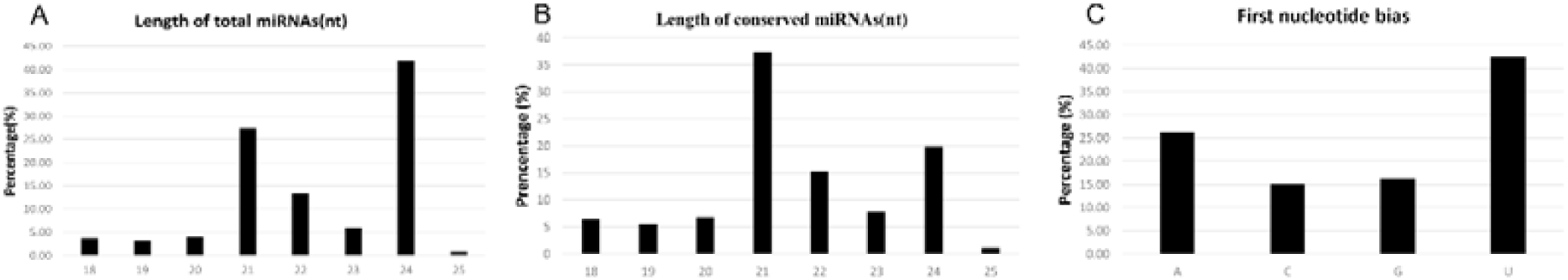
Size distribution of miRNAs and characterization of conversed miRNAs detected by deep sequencing. (A) Length distribution of the total miRNAs; (B) Distribution of obtained unique conversed miRNAs (gp1-gp3); (C) Percentage of first nucleotide bias in the identified conversed miRNAs(gp1-gp3).

### Conserved miRNAs and novel miRNAs identified in Pepper

These unique sequences were subsequently used to identify conserved and novel miRNAs by alignment against miRBase (Version 22), the pepper genome and expressed sequence tag (EST) or pre-miRNA sequences. Unique miRNA transcripts identified from the mappable sequences are divided into five group types (Table S2)(Gong et al., 2015): (1) gp1a type: Unique reads map to specific miRNAs/pre-miRNAs in miRbase and the pre-miRNAs further map to the genome and EST. Total 77 miRNAs from the six samples corresponding to 50 known pepper pre-miRNAs (Table S2-1). (2) gp1a type: Reads map to selected (except for specific) miRNAs/pre-miRNAs in miRbase and the pre-miRNAs further map to the genome and EST. 122 miRNAs, which correspond to 94 known pepper pre-miRNAs, which cannot be mapped to genome or EST (Table S2-2). (3) gp2b type: Unique reads map to selected miRNAs/pre-miRNAs in miRbase. The mapped pre-miRNAs do not map to the genome, but the reads (and of course the miRNAs of the pre-miRNAs) map to genome. The extended genome sequences from the genome loci may form hairpins. 261 miRNAs corresponding to 314 others known miRbase plant pre-miRNAs, which are mapped to genome or EST (Table S2-3). (4) gp3a type: Unique reads map to selected miRNAs/pre-miRNAs in miRbase. The mapped pre-miRNAs do not map to the genome, and the reads do not map to the genome; Forty miRNAs corresponding to 35 others known miRbase plant pre-miRNAs, which cannot be mapped to genome or EST (Table S2-4). (5) gp4a: Unique reads do not map to selected pre-miRNAs in miRBase, but the reads map to genome and the extended genome sequences from genome may form hairpins; 411 miRNAs corresponding to 421 candidate pre-miRNAs, which are predict RNA hairpins derived from genome or EST, and these miRNAs are novel miRNA, which are labeled PC (pepper candidate) (Table S2-5).

In the present study, identified mature miRNAs are divided into two type, including conserved miRNA and novel miRNA (Table 1), referring to the research of Maize(Li et al., 2017). The length of the mature miRNAs ranged from 18 to 25 nt. The 24 nt and 21 nt miRNA category was most abundant, 41.85% and 27.41%, respectively (Figure 2A). This is consistent with the typical lengths of plant sRNAs reported in other studies(Gao et al., 2016;Zhang et al., 2016a). Of the conserved miRNA 21-nt miRNAs were most abundant (37.32%) (Figure 2B), representing the dominant size of mature miRNAs in plants. The 5′ terminal nucleotides of sRNA sequences influence classification of their AGO complexes and is an important feature affecting function(Schuck et al., 2013). Most miRNAs are incorporated into the AGO1 effector complex, resulting in sequence specificity that either cleaves or translationally represses their targets(Dalmadi et al., 2019). Therefore, we examined the 5′ nucleotide distribution of conserved miRNAs, and 42.49% started with uridine at 5’-end, and 26.17% started with adenine (Figure 2C).

### Differentially expressed miRNAs analysis between pepper sterile line N816S and its maintainer line Ning5m

MiRNAs are play an important role in plant development and apoptosis(Jovanovic and Hengartner, 2006;Wang et al., 2007). To determine differential expression between N816S and Ning5m anthers. MiRNA expression was normalized to transcripts per million and simplified as normalized expression (Norm)(Gong et al., 2015). A miRNA was considered if the Norm value was greater than one in the all given replicated samples. Based on this criterion, 525 miRNAs were detected (Figure 3A), including a total of 350 conserved miRNAs (38.42% of the total miRNAs) belonging to 55 families were observed in at least one of the group samples (Table S3). In correlation analysis, Norm values of the N816S and Ning5m anthers were found to be highly correlated between repeats (r>0.99), indicating good reproducibility of the miRNAome results (Figure S1).

**Figure 3.**
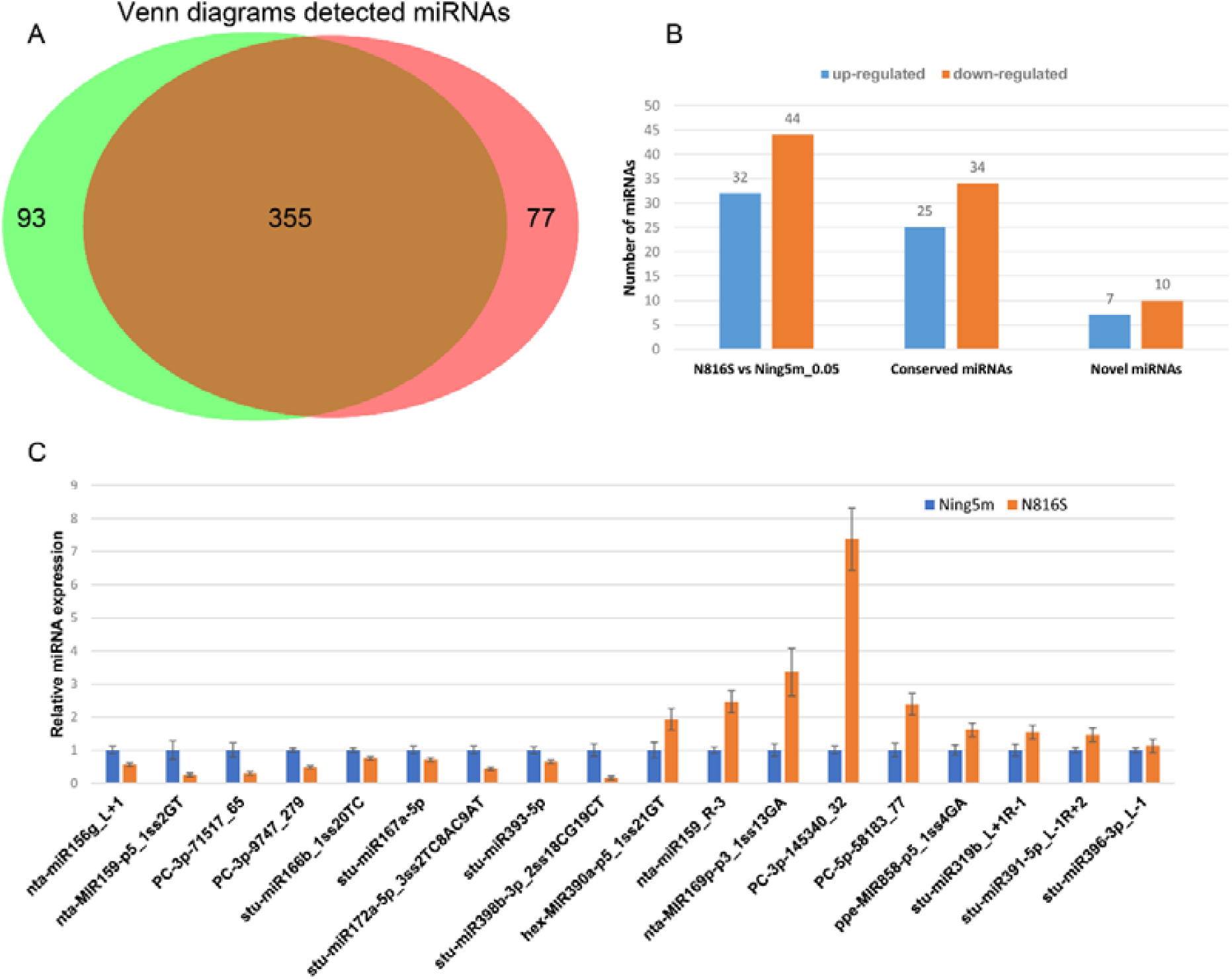
Venn diagrams detected miRNA, the number statistics of differential expression miRNAs, and qRT-PCR for verifying deep sequencing. (A) Detected miRNAs in N816S and Ning5m; (B) The number statistics of up-regulated and down-regulated differential expression miRNAs in conserved miRNAs and novel miRNAs. (C) Detection of selected miRNA expression in N816S and Ning5m anthers using qRT-PCR.

A one-tailed t-test was used to identify differentially expressed miRNAs with p value < 0.05(Xing et al., 2012). The hierarchical clustering of the differential expression mRNAs was made and showed different expression patterns between the N816S and Ning5m (Figure 5A). As a result, a total of 76 miRNAs (59 conserved miRNAs and 17 novel miRNAs) were found to be differentially expressed between the two phenotypes (Table 3). Compared with the Ning5m, 44 miRNAs (34 conserved miRNAs and 10 novel miRNAs) were found to be down-regulated and 32 miRNAs (25 conserved miRNAs and 7 novel miRNAs) up-regulated in the N816S (Figure 3B). Nta-miR156g_L+1 and stu-miR156a-5p had high expression abundance in MiRNA156 family, were down-regulated in sterile line N816S. However, miRNA390 family, including hex-MIR390a-p5_1ss21GT, hex-MIR390b-p5_2ss10TC21GA and hex-MIR390b-p5_1ss21GT, were up-regulated in sterile line N816S. Five miRNAs (stu-miR393-3p, stu-miR399j-3p_1ss21GA, nta-miR6149a_L+1R-1_1ss21GC and stu-miR398b-3p ath-miR8175_L+4) were specifically expressed in Ning5m. Five miRNA(nta-MIR172e-p3_2ss15CG19TA,ppe-MIR399a-p3_2ss5AT18TC,ppe-MIR39 9a-p5_2ss5AT18TC, stu-miR3627-5p_R-1 and sly-MIR10528-p3_2ss9GA19TC) were specifically expressed in N816S. To validate conserved miRNAs identified and novel miRNAs predicted, we selected 18 differential expressed miRNAs for stem-loop qRT-PCR. The expression trends of these miRNAs were consistent with the high-throughput sequencing results. As showed in Figure 3C, expression of these miRNAs from qRT-PCR displayed a similar tendency with those from small RNA sequencing.

**Table 2.** Summary of conserved and predicted miRNA.

| Groups | Pre-miRNA | Unique miRNA | miRNA type |
| --- | --- | --- | --- |
| gp1 | 50 | 77 | Conserved miRNA |
| gp2a | 94 | 122 | Conserved miRNA |
| gp2b | 314 | 261 | Conserved miRNA |
| gp3 | 35 | 40 | Conserved miRNA |
| gp4 | 421 | 411 | Novel miRNA |

**Table 3.**
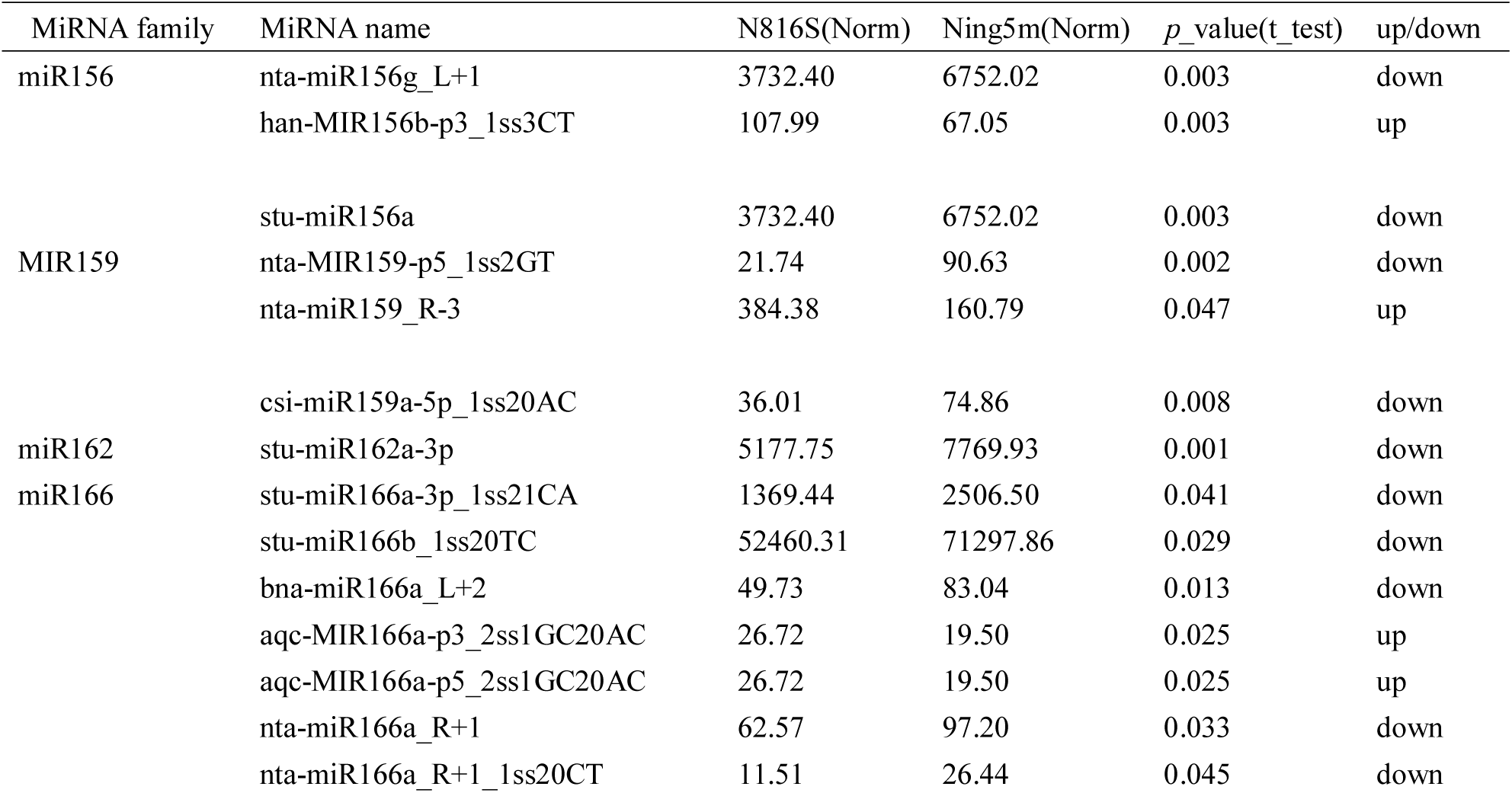
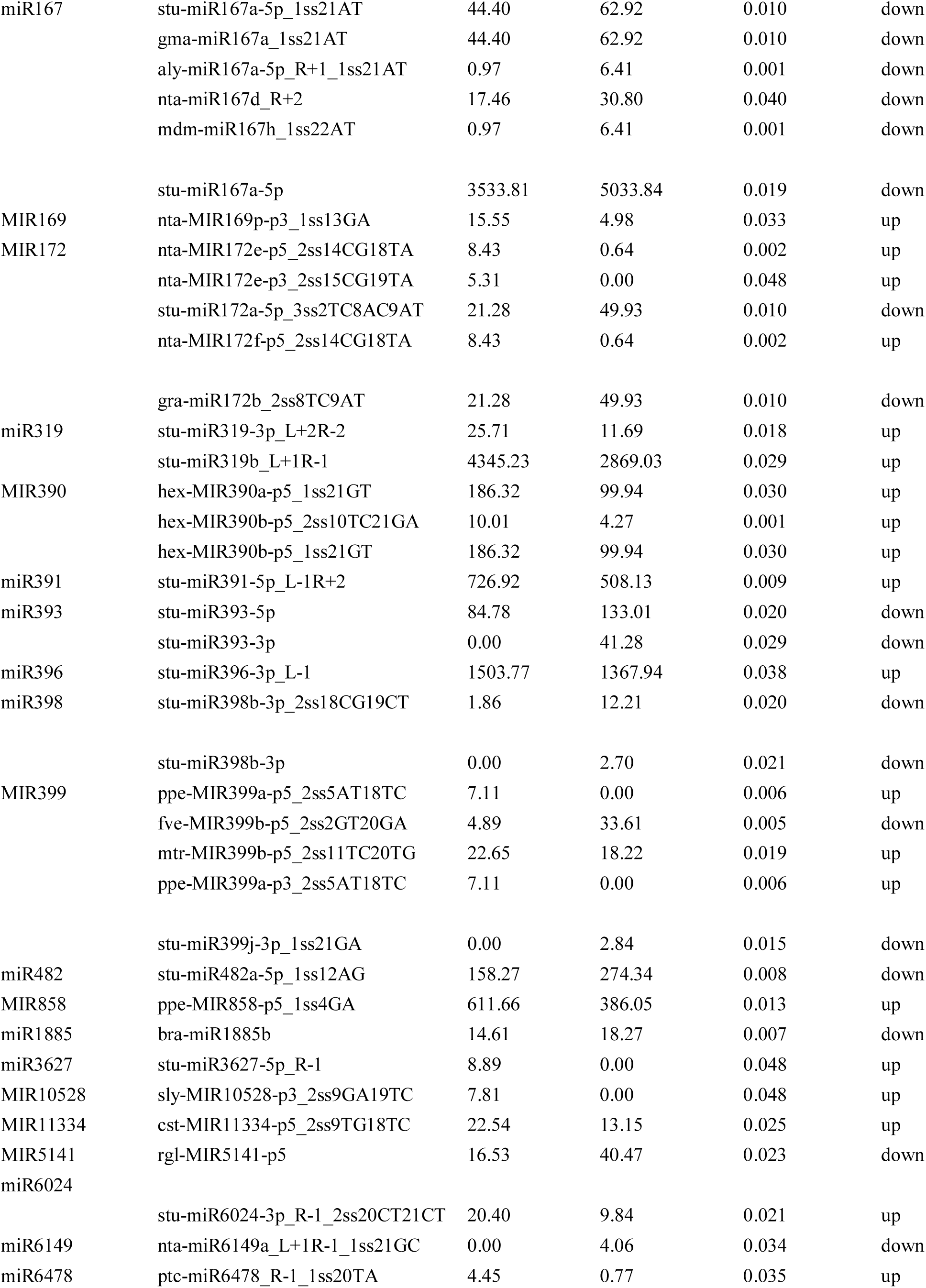
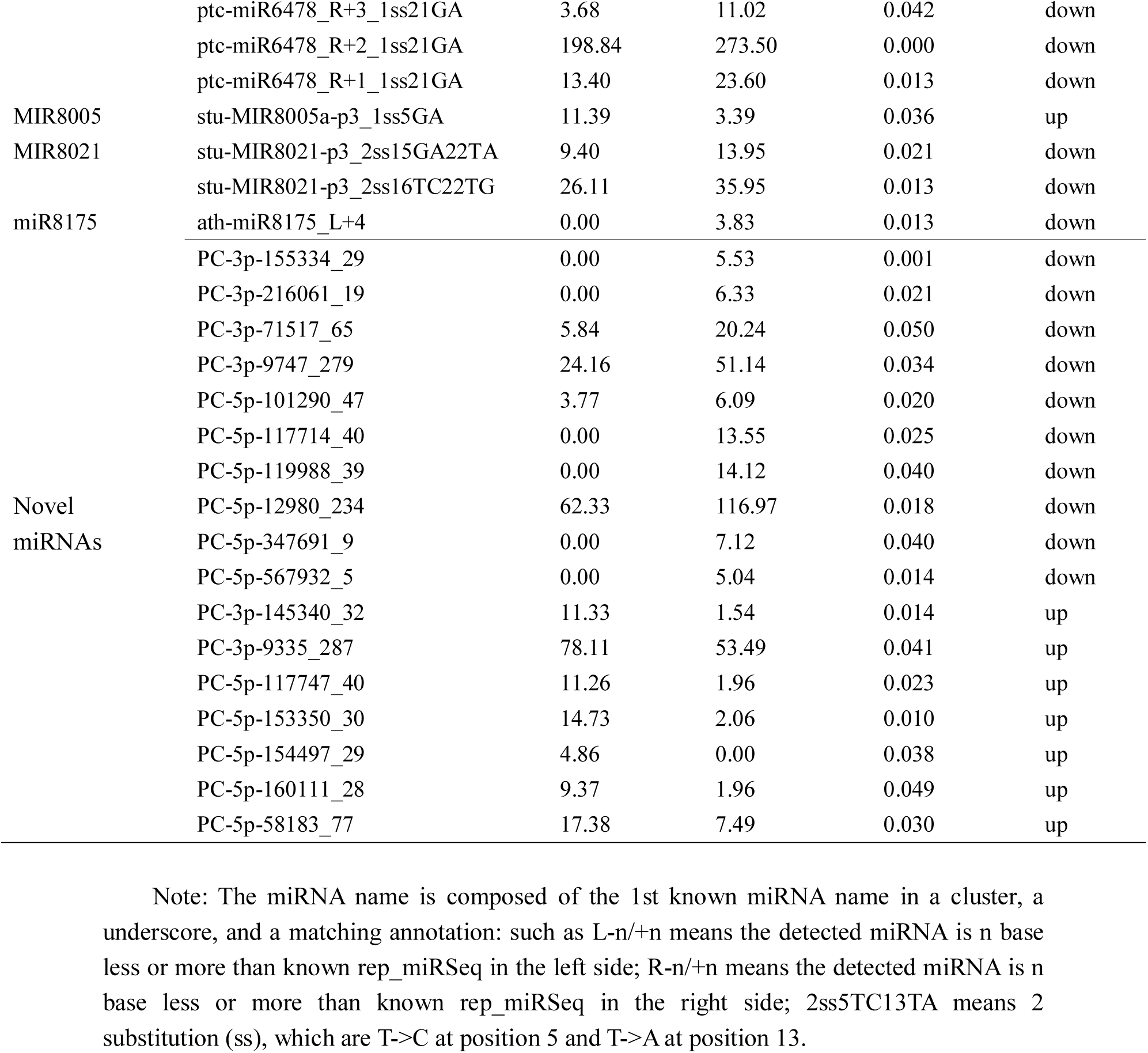
Summary of the differentially expressed conserved and novel miRNAs.

**Figure 4.**
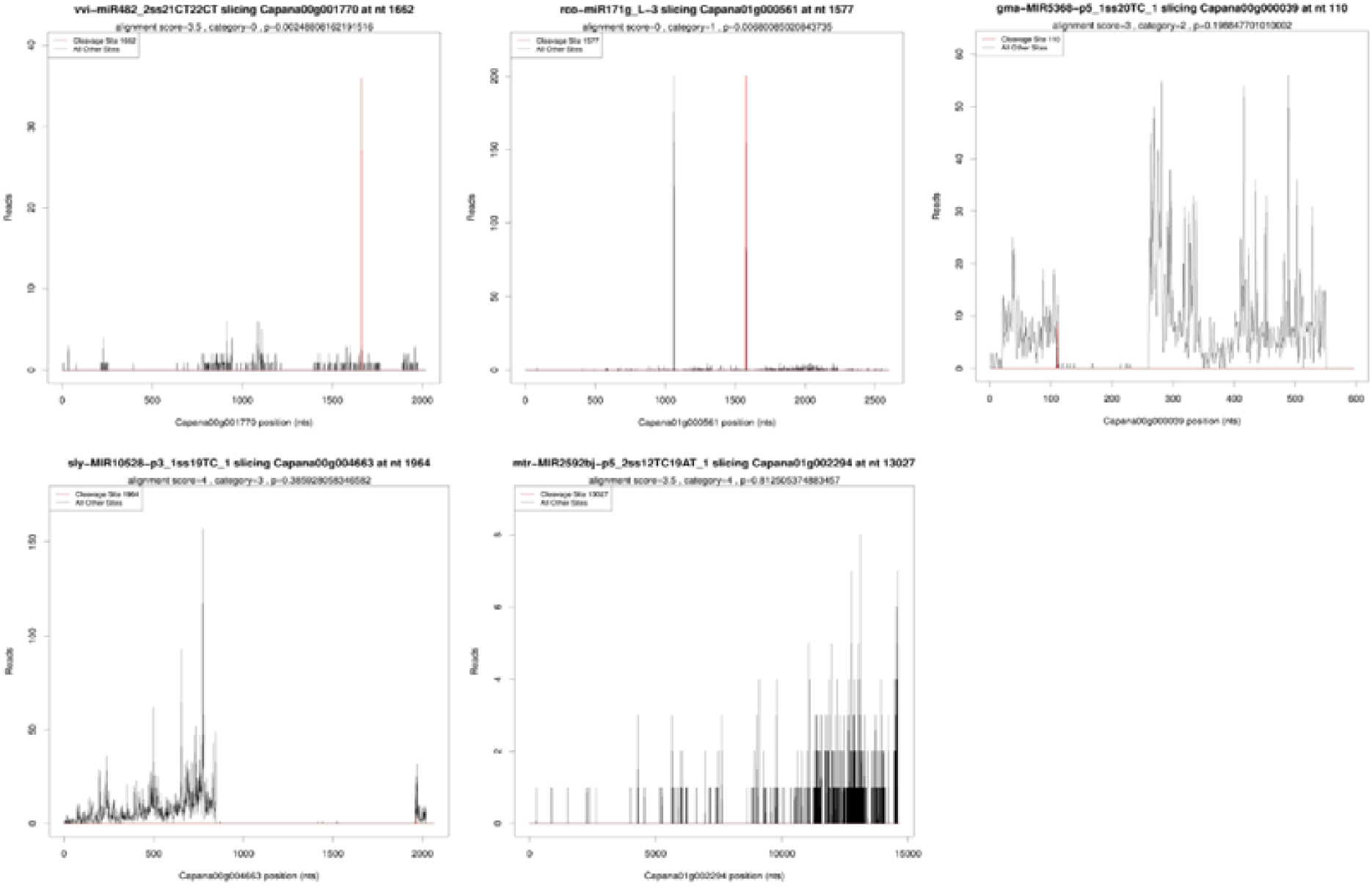
Typical categories of the target transcript according to the relative abundance of the tags at the target mRNA sites.

**Figure 5.**
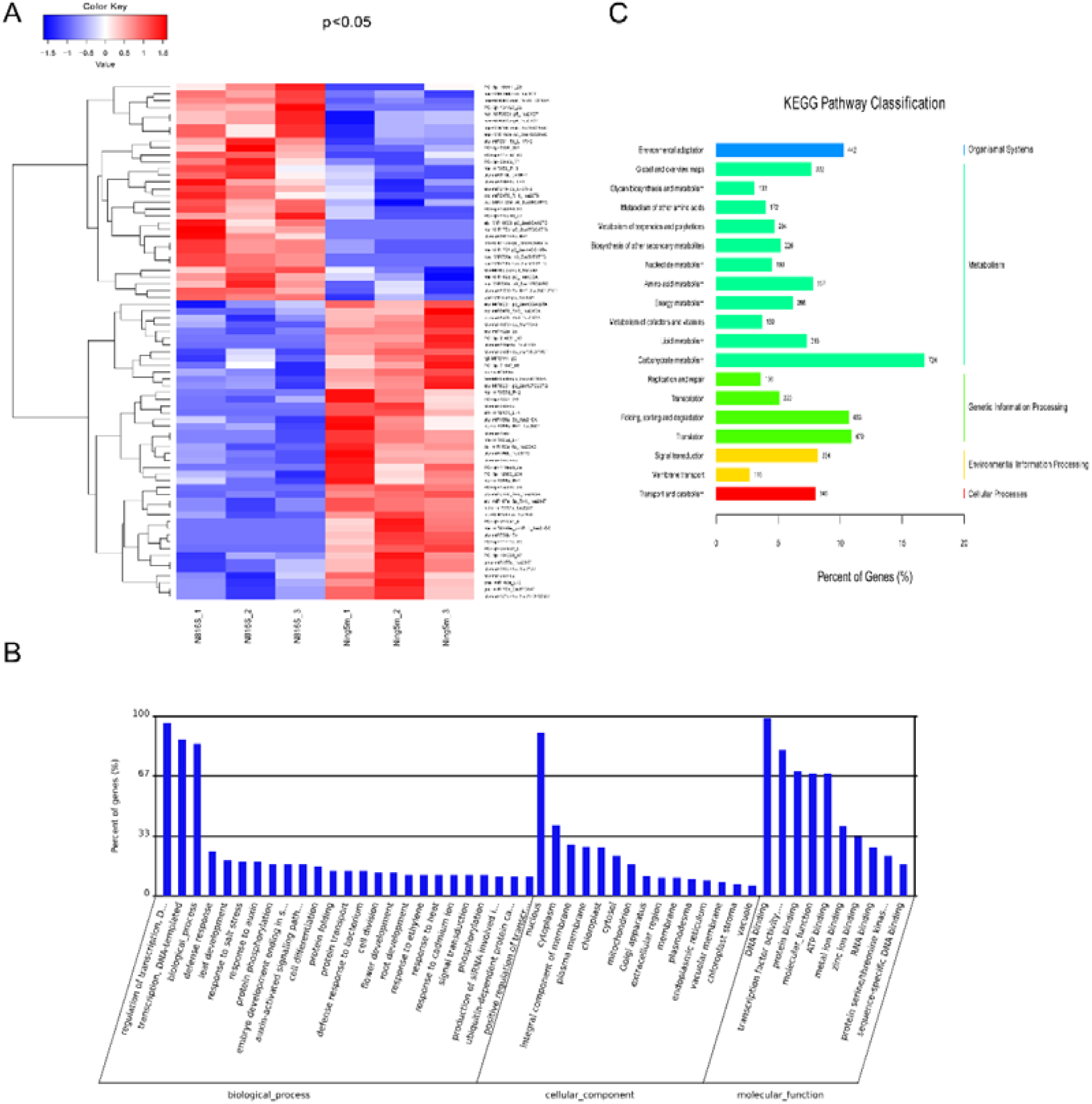
(A) Differentially expressed miRNAs cluster, (B) GO functional classification of miRNA targets, (C) KEGG Pathway Classification; Notes: miRNAs cluster columns are different samples; rows are different miRNAs. Clustering with p_value, red is up-expressed miRNA, blue is down-expressed miRNA.

### Targets analysis of miRNAs in anthers of Pepper

To understand the function of the mature miRNAs from anthers of Pepper, targets of miRNAs were detected using degradome sequencing technology. A total of 34233673 (99.24%) mappable reads from raw reads were obtained, including 12320360(99.19%) unique mappable reads, while 20851831 (60.45%) transcript mapped reads, including 6635426(53.42%) unique transcript mappable reads, were obtained (Table S4). The target transcripts were sorted into five categories according to the relative abundance of the tags at the target mRNA sites(Gong et al., 2015): category “0” is defined as >1 raw read at the position, with abundance at a position equal to the maximum on the transcript and with only one maximum on the transcript; category “1”, the expected cleavage signature was equal to the maximum on the transcript and more than one maximum position on the transcript; category “2”, abundance at the position was less than the maximum but higher than the median for the transcript; category “3”, the abundance at the position equal to or less than the median for the transcript; category “4”, abundance at the position was only one raw read. Figure 6 showed the typical five categories of the target transcripts.

**Figure 6.**
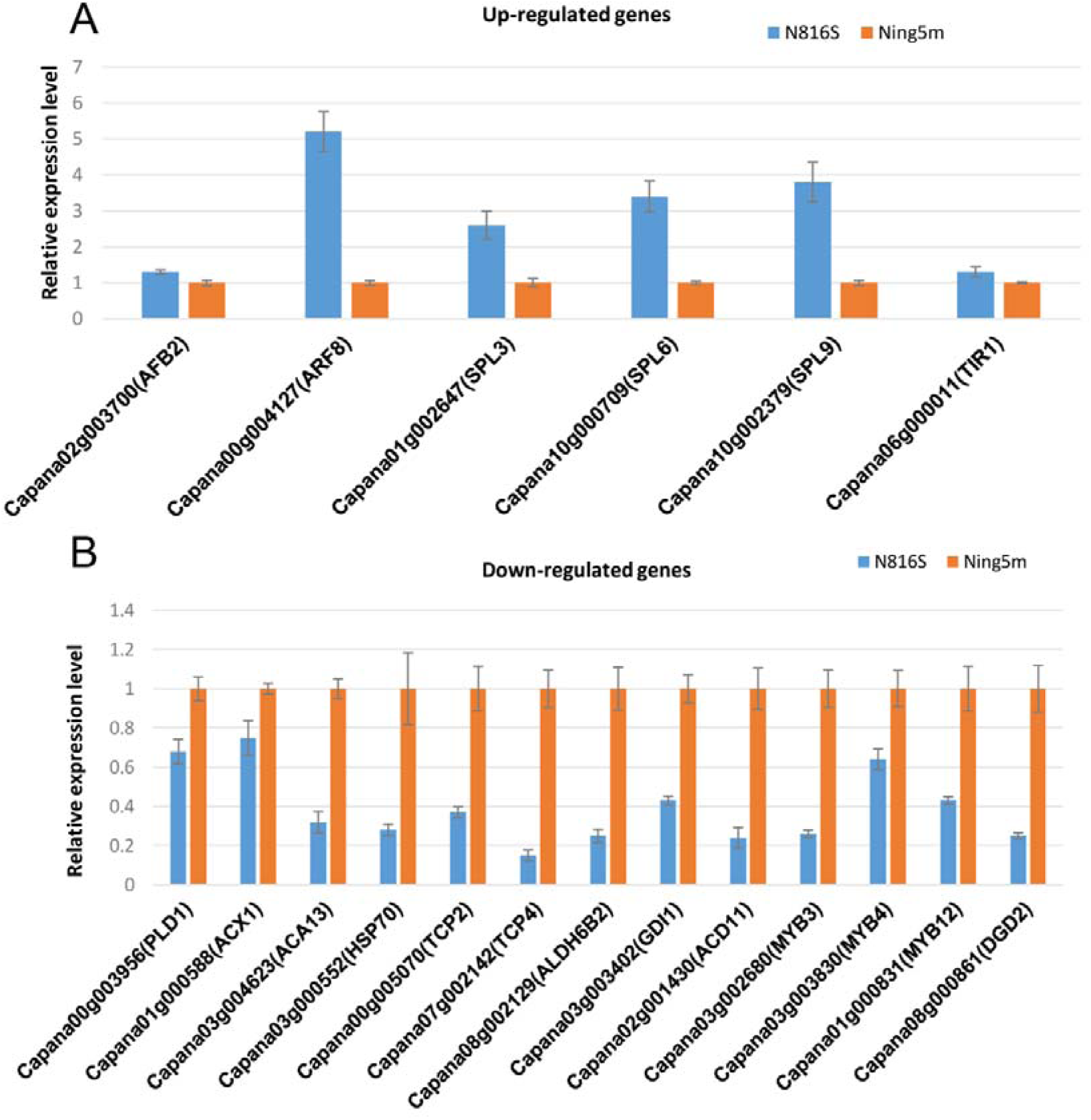
Quantitative real-time PCR analysis of the relative expression of miRNA targets in the CMS-line N816S and its maintainer line Ning5m. (A) up-regulated targets in N816S; (B) down-regulated targets in N816S.

A total of 1292 reliable targets (*p_*value <1) that were potentially cleaved by 321 miRNAs (250 conserved miRNAs and 71 novel miRNAs) were identified (Table S5). A total of 77 target genes were identified to be targeted by 35 differentially expressed miRNAs (27 conserved miRNAs and 8 novel miRNAs) (Table S6). Many targets of miR156, miR167, miRNA858 families were identified as transcription factors, such as SPL, ARF, and MYB, respectively, which have been experimentally validated by the previous studies.

### Gene Ontology (GO) and KEGG Pathway Analysis of Targets

The targets were annotated using the GO annotations analysis, which is commonly used to describe the function of genes and gene products, meanwhile the KEGG analysis is used to provide the pathway of annotated targets. Figure 5B showed the GO functional classification of miRNA targets in the pepper anthers. Targets covered a broad range of functional categories. However, targets of DNA binding, regulation of transcription-, and DNA-templated genes, transcription factor activity-, and sequence-specific DNA binding, transcription-, and DNA-templated genes were mostly enriched. This suggested that those target genes may play an important role in pepper anthers. The pathways of all the miRNAs and targets involved were listed (Table S7). There were 107 pathways that targets involved, while carbohydrate metabolism, translation, folding sorting and degradation pathways owned most targets (Figure 5C). Furthermore, the 77 miRNA targets from 35 differentially expressed miRNAs were found to be involved in multiple pathways, including plant hormone signal transduction, purine metabolism, starch and sucrose metabolism, oxidative phosphorylation, and others (Table S8).

### Expression profiles of miRNA targets examined by qRT-PCR

To examine the correlation between the targets and the corresponding miRNAs, the expression levels of 19 selected targets were examined by qRT-PCR analysis. MiRNAome sequencing result showed that miRNA156a/g (stu-miR156a and nta-miR156g_L+1), miRNA167a/h (stu-miR167a-5p, stu-miR167a-5p_1ss21AT and mdm-miR167h_1ss22AT), stu-miR393-5p and nta-miR6149a_L+1R-1_1ss21GC were down-regulated in N816S compare to Ning5m (Table 3). The degradome sequencing results indicate that miRNA156a/g target SPL family genes (SPL3, SPL6 and SPL9), miRNA167a/h (aly-miR167a-5p_R+1_1ss21AT, gma-miR167a_1ss21AT, mdm-miR167h_1ss22AT, stu-miR167a-5p_1ss21AT) targets an AUXIN RESPONSE FACTOR gene (ARF8), MiRNA393 targets two AUXIN SIGNALING F□BOX genes (TIR1 and AFB2) (Table S8). QRT-PCR analysis showed that the expression profiles of the above miRNA targets were up-regulated in N816S (Figure 6A).

In contrast, miRNA159_R3, nta-MIR169j-p3_2ss8GA19CT, stu-miR319b_L+1R-1, hex-MIR390a-p5_1ss21GT, ppe-MIR399a-p3_2ss5AT18TC, ppe-MIR858-p5_1ss4GA, cst-MIR11334-p5_2ss9TG18TC and PC-5p-154497_29 were up-regulated in N816S (Table 3). MiRNA159_R3 targets a plasma membrane H+-ATPase gene (*ACA13*), a γ-TuC Protein 3 (GCP3)-Interacting Protein gene (*GIP1*), MiRNA169j-p3 targets a heat shock protein gene (HSP70), MiRNA319b targets four TEOSINTE BRANCHED/CYCLOIDEA/PCF transcription factor genes (*TCP2, TCP4, TCP24*) and a Aldehyde dehydrogenase gene (*ALDH6B2*), MiRNA390a-p5 targets a GDP dissociation inhibitor gene (*GDI1*), miRNA399a-p3 targets a ceramide-1-phosphate transfer protein gene (*ACD11*), miRNA858-p5 targets several MYB protein genes (*MYB3, MYB4, MYB12* and *MYB80*), miR11334-p5 targets a phospholipase D alpha 1-like gene (*PLD1*) and a Acyl-CoA oxidase gene (*ACX1*), and a novel miRNA of PC-5p-154497_29 targets a UDP-galactose-dependent digalactosyldiacylglycerol synthase gene (*DGD2*) (Table S8). Most of above targets were down-regulated in N816S (Figure 6B).

Most negative correlations were found between the expression levels of the target genes and their corresponding miRNAs in anthers of the sterile line N816S and its maintainer line Ning5m, implying that miRNA-mediated mRNA silencing occurred during anther development.

## Discussion

Several studies have shown that miRNAs regulate anther development in plants(Yang et al., 2013;Gong et al., 2015). However, few studies on the relationships between miRNA biogenesis and CMS occurrence during anther development in pepper were very limited. MiRNAome and degradation sequencing technology provides an effective way to identify and evaluate the expression profiles of miRNAs and targets associated with CMS occurrence in plant tissues during anther development(Wei et al., 2015;Hu et al., 2016;Bai et al., 2017). To explore the roles of miRNAs during the occurrence of CMS, anthers at the early uninucleate stage were used to examine the expression profiles of the miRNAs in the sterile line N816S and its maintainer line Ning5m using high-throughput sequencing. To the best of our current knowledge, this study is the first report on identification and characterization of miRNAs, and their targets between a sterile line and its maintainer line during anther development in pepper.

In this study, a total of 76 miRNAs (59 conserved miRNAs and 17 novel miRNAs) were identified as differentially expressed between the sterile line N816S and its maintainer line Ning5m. To understand the function of the miRNAs from anthers of Pepper, targets of miRNAs were detected using degradome sequencing technology. From those targets, 80 target genes were identified to be targeted by 35 differentially expressed miRNAs (27 conserved miRNAs and 8 novel miRNAs). The qRT-PCR results indicated that most negative correlations were found between the expression levels of the target genes and their corresponding miRNAs in the anthers of the sterile line N816S and its maintainer line Ning5m. KEGG analysis showed that most gene targets of the 35 differentially expressed miRNAs were involved in pathways, including plant hormone signal transduction, starch and sucrose metabolism, oxidative phosphorylation, purine metabolism, and others.

Intriguingly, a number of genes in hormone signaling were confirmed or predicted as targets of miRNAs. Auxin regulates anther dehiscence, pollen maturation, and filament elongation in *Arabidopsis*. The miR167-guided cleavage of auxin response factor *ARF8*, miR393-guided cleavage of auxin receptor *TIR1* and *AFB2*, and miR319-guided cleavage of jasmone acid biosynthesis related transcription factor genes *TCPs*, which were reported in previous study(Liu and Chen, 2009). In *Arabidopsis*, loss of miR167 regulation in *mARF6* and *mARF8* expression caused arrested ovule development and anther indehiscence(Wu et al., 2006). Four auxin receptor-encoding genes, *TIR1, AFB1, AFB2*, and *AFB3*, are transcribed in anthers only during late stages of development starting at the end of meiosis(Shimizu-Mitao and Kakimoto, 2014). Up-regulation of *TIR1* enhances auxin sensitivity, and causes altered leave phenotype and delayed flowering in *Arabidopsis*(Chen et al., 2011). MicroRNA319-regulated TCPs can activate CO transcription and control flowering time, shaping flower structure, leaf morphology, and plant architecture in Arabidopsis(Chen et al., 2011). In this study, up-regulation *AFR8* was targeted by down-regulated miRNA393a/h, up-regulated *TIR1* and *AFB2* were targeted by down-regulated miR393a, and down-regulated *TCPs* (*TCP2* and *TCP4*) were targeted by up-regulated miRNA393b, indicating might modulate the hormone response to play roles in the pollen development and CMS occurrence.

Targets of these differentially expressed miRNAs containing important transcription factors (TFs) and functional proteins are involved in many biological processes, including signal transduction, floral organ development, and organellar gene expression(Liu and Chen, 2009). For instance, miR156 targets *SPL* transcription factor family, which is regulatory functions throughout the growth and development stages in plants. Previous report showed that miR156 regulates the timing of flower formation vis *SPL3/4/5*, activating the expression of *LEAFY, FRUITFULL* and *APETALA*(Jung et al., 2011). In *Arabidopsis*, multiple *SPL* genes regulate cell division, differentiation and can result in fertile flower(Zheng et al., 2019). In this study, *SPL3, SPL6* and *SPL9* were identified as potentially targets of miR156 miRNA family members (stu-miR156a, stu-miR156a, nta-miR156g_L+1) were identified as approximately triple up-regulated in N816S compared with Ning5M (Figure 6A), leading to disordered floral organ development in pepper, indicating that miR156 may participate in fertility regulation. Furthermore, miR858 targets *MYB* transcription factor family, which is involved in the control of plant development, determination of cell fate and identity, primary, and secondary metabolism. In rice, anther and pollen defect in floral organ development are found in the loss-of-function mutations of *MYB*. In Arabidopsis, *AtMYB3, AtMYB4, AtMYB7* and *AtMYB32* encode transcriptional repressors, and *AtMYB4* controls sinapate ester biosynthesis, whereas *AtMYB32* regulates pollen wall composition, and AtMYB12 is involved in the regulation of flavonoid biosynthesis control flavonol biosynthesis(Dubos et al., 2010). The transcriptional activity of the PtMYB4 protein is positively regulated by the mitogen-activated protein kinase (MAPK) PtMAPK6 in *Pinus taeda*, which phosphorylates a Ser in the C terminal activation domain. In addition, *AtMYB80* regulated exine formation and acts downstream of *AtMYB35* to control anther development and/ or functionalities. In this present study, up-regulated miRNA858 might via reducing transcript expression of *MYB3, MYB4, MYB12* and *MYB80*, leading to anther and pollen defect. In addition, MIR169j targets a heat shock protein *HSP70*, which has been widely involved in the protein peptides folding, assembly and transports, and the degradation of abnormal proteins. Studies have found that *HSP70* is associated with male sterility in plant and animal. Down regulation of *HSP70* expression level prolongs the duration of heat□induced male sterility in *Drosophila buzzatii*(Sarup et al., 2004). *HSP70* antisense RNA expression leads to male sterility in rice(Liu et al., 2008). In this study, up-regulated miRNA169j may through via reducing transcript expression of *HSP70*, leading to pollen abortion.

Many miRNA-targeted genes were involved in lipid transport and metabolism, such as *PLD1/ACX1, ACA13/GIP, ALDH6, GDI1* and *DGD2*, which are cleaved by miR11334, miR159_R-3, miR319b, miRNA390b and PC-5p-154497_29, respectively. Those miRNAs may throughout their targets be involved in CMS sterile process. Phospholipase D alpha 1-like (PLD1) has been identified as cytosolic protein, which regulated cytosolic lipid droplet formation(Andersson et al., 2006). Acyl-CoA oxidase (ACX) was the first and the key step controlling enzyme involved in fatty acid β-oxidation, and mutation of *ACX1* leaded to petal degeneration in Chinese Cabbage(Zheng et al., 2019). In plant, *ALDH6B2* encodes a methylmalonyl semialdehyde dehydrogenase, of which involved in the degradation of valine to propionyl CoA(Brocker et al., 2013). In addition, Plant cells have multiple plasma membrane (PM)-localized calcium ATPases (ACAs) pumping calcium ions out of the cytosol. The loss of ACA13 combination with a reduction in function of other ACAs leads to seedling death at bolting, revealing the essential role of their collective function in plant growth(Yu et al., 2018). Moreover, the γ-tubulin complex (γ-TuC) Protein 3 (GCP3)-Interacting Protein 1 (GIP1) is the smallest γ-TuC component identified. In Arabidopsis, *AtGIP1* and its homologous protein *AtGIP2* mutants are impaired in establishing a fully functional mitotic spindle and exhibit severe developmental defects(Batzenschlager et al., 2013). The GDP dissociation inhibitor protein GDI1 relates to control vesicle number and transport in an amelioration of zinc toxicity, allowing yeast to survive in the presence of toxic(Ezaki and Nakakihara, 2012). Furthermore, Digalactosyl-diacylglycerol (DGD) is one of the major lipids found predominantly in the photosynthetic membrane of higher plants. *OsDGD2*β is the sole DGDG synthase gene highly expressed in anther, and its mutation confers male sterility with pale yellow and shrunken anther, devoid of starch granules in pollen, and delayed degeneration of tapetal cells in rice(Basnet et al., 2019). All above related miRNAs are up-regulated in sterile line N816S compare to its maintainer Ning5m and those disorder miRNA-targeted genes perhaps leading to membrane-disruptive effects and cell apoptosis.

## Conclusion

In the present study, small RNA libraries from anther of CMS-line N816S and its maintainer line Ning5m were generated by small RNA sequencing in pepper. A total of 76 differentially expressed miRNAs were discovered, of which 18 were further confirmed by real-time quantitative PCR (qRT-PCR). Furthermore, targets of miRNAs were identified by degradome sequencing. A total of 1292 targets that were potentially cleaved by 321 miRNAs (250 conserved miRNAs and 71 novel miRNAs) were identified. Gene Ontology (GO) and KEGG pathway analysis of target transcripts indicated that 77 target genes cleaved by 35 differentially expressed miRNAs might play roles in the regulation of CMS sterility, such as MYB, SPL, and AFR family proteins targeted by miR156, miR167, miRNA858 family. Nineteen targets were selectively examined by qRT-PCR, and the results showed that there was a negative correlation on the expression patterns between miRNAs and their targets. These findings provide valuable information to understand the roles of miRNAs during anther development and CMS occurrence in pepper.

## Author contributions

Min Zhang designed the study. Hongyuan Zhang, Shuping Huang and Jie Tan carried out the experiment, data analysis, interpretation of the results. Hongyuan Zhang drafted the manuscript. Xia Chen supervised the work and revised the manuscript. All authors have read and approved the final version of this submission.

## Acknowledgement

This work was financially supported by the National Natural Science Foundation of China (Grant No. 31701936) and the subproject of National key research and development Program of China (2017YFD0101904).

## Supporting Information

Figure S1. Small RNAs Pearson correlation between the six libraries

Table S1. Primers used in this study. S1-1, Primers designed for conserved miRNAs and novel miRNAs; S1-2 Primers designed for miRNA targets.

Table S2. Summary of five types of miRNA in this study. S2-1, gp1a type; S2-2, gp2a type; S2-3, gp2b type; S2-4, gp3 type; S2-5, gp4 type.

Table S3. Primers used in this study. S1-1, Primers designed for conserved miRNAs and novel miRNAs; S1-2, Primers designed for miRNA targets.

Table S4. Overview of degradome sequencing reads from raw data to mapping sequences.

Table S5. The overview of reliable identified targets of miRNAs.

Table S6. Transcript annotation of targets of differential expression conserved and novel miRNAs in this study.

Table S7. The pathways of all the miRNAs and targets involved in this study.

Table S8. The transcript annotation, GO terms and KEGG pathways of differential expression miRNAs with those targets in pepper.

## Reference

Andersson, L., Bostrom, P., Ericson, J., Rutberg, M., Magnusson, B., Marchesan, D., Ruiz, M., Asp, L., Huang, P., Frohman, M.A., Boren, J., and Olofsson, S.O. (2006). PLD1 and ERK2 regulate cytosolic lipid droplet formation. J Cell Sci 119, 2246–2257.

Anthony, A.M.F. G. (2005). The Arabidopsis GAMYB-Like Genes, MYB33 and MYB65, Are MicroRNA-Regulated Genes That Redundantly Facilitate Anther Development. The Plant Cell 17, 705–721.

Asha, S., Sreekumar, S., and Soniya, E.V. (2016). Unravelling the complexity of microRNA-mediated gene regulation in black pepper (Piper nigrum L.) using high-throughput small RNA profiling. Plant Cell Reports 35, 53–63.

Bai, J.F., Wang, Y.K., Wang, P., Duan, W.J., Yuan, S.H., Sun, H., Yuan, G.L., Ma, J.X., Wang, N., Zhang, F.T., Zhang, L.P., and Zhao, C.P. (2017). Uncovering Male Fertility Transition Responsive miRNA in a Wheat Photo-Thermosensitive Genic Male Sterile Line by Deep Sequencing and Degradome Analysis. Frontiers In Plant Science 8.

Barrajon-Catalan, E., Alvarez-Martinez, F.J., Borras, F., Perez, D., Herrero, N., Ruiz, J.J., and Micol, V. (2020). Metabolomic analysis of the effects of a commercial complex biostimulant on pepper crops. Food Chemistry 310.

Basnet, R., Hussain, N., and Shu, Q. (2019). OsDGD2β is the Sole Digalactosyldiacylglycerol Synthase Gene Highly Expressed in Anther, and its Mutation Confers Male Sterility in Rice. Rice 12, 66.

Batzenschlager, M., Masoud, K., Janski, N., Houlné, G., Herzog, E., Evrard, J.-L., Baumberger, N., Ehrardt, M., Nominé, Y., Kieffer, B., Schmit, A.-C., and Chabouté, M.-E. (2013). The GIP gamma-tubulin complex-associated proteins are involved in nuclear architecture in Arabidopsis thaliana. Frontiers in Plant Science 4.

Bohra, A., Jha, U.C., Adhimoolam, P., Bisht, D., and Singh, N.P. (2016). Cytoplasmic male sterility (CMS) in hybrid breeding in field crops. Plant Cell Reports 35, 967–993.

Brocker, C., Vasiliou, M., Carpenter, S., Carpenter, C., Zhang, Y., Wang, X., Kotchoni, S.O., Wood, A.J., Kirch, H.-H., Kopečný, D., Nebert, D.W., and Vasiliou, V. (2013). Aldehyde dehydrogenase (ALDH) superfamily in plants: gene nomenclature and comparative genomics. Planta 237, 189–210.

Chen, Z.H., Bao, M.L., Sun, Y.Z., Yang, Y.J., Xu, X.H., Wang, J.H., Han, N., Bian, H.W., and Zhu, M.Y. (2011). Regulation of auxin response by miR393-targeted transport inhibitor response protein 1 is involved in normal development in Arabidopsis. Plant Molecular Biology 77, 619–629.

Dalmadi, A., Gyula, P., Balint, J., Szittya, G., and Havelda, Z. (2019). AGO-unbound cytosolic pool of mature miRNAs in plant cells reveals a novel regulatory step at AGO1 loading. Nucleic Acids Research 47, 9803–9817.

Diaz-Manzano, F.E., Cabrera, J., Ripoll, J.J., Del Olmo, I., Andres, M.F., Silva, A.C., Barcala, M., Sanchez, M., Ruiz-Ferrer, V., De Almeida-Engler, J., Yanofsky, M.F., Pineiro, M., Jarillo, J.A., Fenoll, C., and Escobar, C. (2018). A role for the gene regulatory module microRNA172/TARGET OF EARLY ACTIVATION TAGGED 1/FLOWERING LOCUS T (miRNA172/TOE1/FT) in the feeding sites induced by Meloidogyne javanica in Arabidopsis thaliana. New Phytologist 217, 813–827.

Dubos, C., Stracke, R., Grotewold, E., Weisshaar, B., Martin, C., and Lepiniec, L. (2010). MYB transcription factors in Arabidopsis. Trends in Plant Science 15, 573–581.

Ezaki, B., and Nakakihara, E. (2012). Possible involvement of GDI1 protein, a GDP dissociation inhibitor related to vesicle transport, in an amelioration of zinc toxicity in Saccharomyces cerevisiae. Yeast 29, 17–24.

Gao, S., Yang, L., Zeng, H.Q., Zhou, Z.S., Yang, Z.M., Li, H., Sun, D., Xie, F.L., and Zhang, B.H. (2016). A cotton miRNA is involved in regulation of plant response to salt stress. Scientific Reports 6.

German, M.A., Luo, S.J., Schroth, G., Meyers, B.C., and Green, P.J. (2009). Construction of Parallel Analysis of RNA Ends (PARE) libraries for the study of cleaved miRNA targets and the RNA degradome. Nature Protocols 4, 356–362.

Gong, S.M., Ding, Y.F., Huang, S.X., and Zhu, C. (2015). Identification of miRNAs and Their Target Genes Associated with Sweet Corn Seed Vigor by Combined Small RNA and Degradome Sequencing. Journal Of Agricultural And Food Chemistry 63, 5485–5491.

Hu, J.H., Jin, J., Qian, Q., Huang, K.K., and Ding, Y. (2016). Small RNA and degradome profiling reveals miRNA regulation in the seed germination of ancient eudicot Nelumbo nucifera. Bmc Genomics 17.

Huang, Q.X., Cheng, X.Y., Mao, Z.C., Wang, Y.S., Zhao, L.L., Yan, X., Ferris, V.R., Xu, R.M., and Xie, B.Y. (2010). MicroRNA Discovery and Analysis of Pinewood Nematode Bursaphelenchus xylophilus by Deep Sequencing. Plos One 5.

Hwang, D.G., Park, J.H., Lim, J.Y., Kim, D., Choi, Y., Kim, S., Reeves, G., Yeom, S.I., Lee, J.S., Park, M., Kim, S., Choi, I.Y., Choi, D., and Shin, C. (2013). The Hot Pepper (Capsicum annuum) MicroRNA Transcriptome Reveals Novel and Conserved Targets: A Foundation for Understanding MicroRNA Functional Roles in Hot Pepper. Plos One 8.

Jeyaraj, A., Wang, X.W., Wang, S.S., Liu, S.R., Zhang, R., Wu, and Wei, C.T. (2019). Identification of Regulatory Networks of MicroRNAs and Their Targets in Response to Colletotrichum gloeosporioides in Tea Plant (Camellia sinensis L.). Frontiers In Plant Science 10.

Jha, A., and Shankar, R. (2011). Employing machine learning for reliable miRNA target identification in plants. Bmc Genomics 12.

Jifon, J., Crosby, K., Patil, B., and Leskovar, D. (2019). Effects of Potassium Nutrition on Seed Yield and Quality of Hybrid Chili Pepper (Capsicum annuum L.). Hortscience 54, S395–S395.

Jovanovic, M., and Hengartner, M.O. (2006). miRNAs and apoptosis: RNAs to die for. Oncogene 25, 6176–6187.

Jung, J.-H., Seo, P.J., Kang, S.K., and Park, C.-M. (2011). miR172 signals are incorporated into the miR156 signaling pathway at the SPL3/4/5 genes in Arabidopsis developmental transitions. Plant Molecular Biology 76, 35–45.

Kim, D.H., Kang, J.G., and Kim, B.D. (2007). Isolation and characterization of the cytoplasmic male sterility-associated orf456 gene of chili pepper (Capsicum annuum L.). Plant Mol Biol 63, 519–532.

Li, H.P., Peng, T., Wang, Q., Wu, Y.F., Chang, J.F., Zhang, M.B., Tang, G.L., and Li, C.H. (2017). Development of Incompletely Fused Carpels in Maize Ovary Revealed by miRNA, Target Gene and Phytohormone Analysis. Frontiers In Plant Science 8.

Lin, C.S., Chen, J.J.W., Huang, Y.T., Hsu, C.T., Lu, H.C., Chou, M.L., Chen, L.C., Ou, C.I., Liao, D.C., Yeh, Y.Y., Chang, S.B., Shen, S.C., Wu, F.H., Shih, M.C., and Chan, M.T. (2013). Catalog of Erycina pusilla miRNA and categorization of reproductive phase-related miRNAs and their target gene families. Plant Molecular Biology 82, 193–204.

Liu, L., Liu, G.Q., and Hou, N. (2008). Construction and identification of HSP70 antisense RNA expression vector for genetic engineering male sterility in plant. Anhui Agricultural Science and Technology 9, 84, 128.

Liu, Q., and Chen, Y.Q. (2009). Insights into the mechanism of plant development: Interactions of miRNAs pathway with phytohormone response. Biochemical And Biophysical Research Communications 384, 1–5.

Parra-Vega, V., Gonzalez-Garcia, B., and Segui-Simarro, J.M. (2013). Morphological markers to correlate bud and anther development with microsporogenesis and microgametogenesis in pepper (Capsicum annuum L.). Acta Physiologiae Plantarum 35, 627–633.

Pei, H.X., Ma, N., Chen, J.W., Zheng, Y., Tian, J., Li, J., Zhang, S., Fei, Z.J., and Gao, J.P. (2013). Integrative Analysis of miRNA and mRNA Profiles in Response to Ethylene in Rose Petals during Flower Opening. Plos One 8.

Reuter, J., and Mathews, D.H. (2009). RNAstructure: Software for RNA Secondary Structure Prediction and Analysis. Journal Of Biomolecular Structure & Dynamics 26, 831–832.

Ru, P., Xu, L., Ma, H., and Huang, H. (2006). Plant fertility defects induced by the enhanced expression of microRNA167. Cell Research 16, 457–465.

Sarup, P., Dahlgaard, J., Norup, A.M., Jorgensen, K.T., Hebsgaard, M.B., and Loschcke, V. (2004). Down regulation of Hsp70 expression level prolongs the duration of heat-induced male sterility in Drosophila buzzatii. Functional Ecology 18, 365–370.

Schuck, J., Gursinsky, T., Pantaleo, V., Burgyan, J., and Behrens, S.E. (2013). AGO/RISC-mediated antiviral RNA silencing in a plant in vitro system. Nucleic Acids Research 41, 5090–5103.

Shimizu-Mitao, Y., and Kakimoto, T. (2014). Auxin Sensitivities of All Arabidopsis Aux/IAAs for Degradation in the Presence of Every TIR1/AFB. Plant And Cell Physiology 55, 1450–1459.

Stepien, A., Knop, K., Dolata, J., Taube, M., Bajczyk, M., Barciszewska-Pacak, M., Pacak, A., Jarmolowski, A., and Szweykowska-Kulinska, Z. (2017). Posttranscriptional coordination of splicing and miRNA biogenesis in plants. Wiley Interdisciplinary Reviews-Rna 8.

Tripathi, R.K., Bregitzer, P., and Singh, J. (2018). Genome-wide analysis of the SPL/miR156 module and its interaction with the AP2/miR172 unit in barley. Scientific Reports 8.

Wang, Y., Stricker, H.M., Gou, D., and Liu, L. (2007). MicroRNA: past and present. Front Biosci 12, 2316–2329.

Wei, X.C., Zhang, X.H., Yao, Q.J., Yuan, Y.X., Li, X.X., Wei, F., Zhao, Y.Y., Zhang, Q., Wang, Z.Y., Jiang, W.S., and Zhang, X.W. (2015). The miRNAs and their regulatory networks responsible for pollen abortion in Ogura-CMS Chinese cabbage revealed by high-throughput sequencing of miRNAs, degradomes, and transcriptomes. Frontiers In Plant Science 6.

Wu, M.F., Tian, Q., and Reed, J.W. (2006). Arabidopsis microRNA167 controls patterns of ARF6 and ARF8 expression, and regulates both female and male reproduction. Development 133, 4211–4218.

Xing, R.H., Lussier, Y.A., Salama, J.K., Khodarev, N.N., Huang, Y., Zhang, Q.B., Khan, S.A., Yang, X.N., Hasselle, M.D., Darga, T.E., Malik, R., Fan, H.L., Perakis, S., Filippo, M., Corbin, K., Lee, Y., Posner, M.C., Chmura, S.J., Hellman, S., and Weichselbaum, R.R. (2012). MicroRNA expression characterizes oligometastasis(es). Cancer Research 72.

Yang, J., Liu, X., Xu, B., Zhao, N., Yang, X., and Zhang, M. (2013). Identification of miRNAs and their targets using high-throughput sequencing and degradome analysis in cytoplasmic male-sterile and its maintainer fertile lines of Brassica juncea. BMC Genomics 14, 9.

Yeri, A., Courtright, A., Danielson, K., Hutchins, E., Alsop, E., Carlson, E., Hsieh, M., Ziegler, O., Das, A., Shah, R.V., Rozowsky, J., Das, S., and Van Keuren-Jensen, K. (2018). Evaluation of commercially available small RNASeq library preparation kits using low input RNA. Bmc Genomics 19.

Yu, H., Yan, J., Du, X., and Hua, J. (2018). Overlapping and differential roles of plasma membrane calcium ATPases in Arabidopsis growth and environmental responses. J Exp Bot 69, 2693–2703.

Yu, S., Galvao, V.C., Zhang, Y.C., Horrer, D., Zhang, T.Q., Hao, Y.H., Feng, Y.Q., Wang, S., Schmid, M., and Wang, J.W. (2012). Gibberellin Regulates the Arabidopsis Floral Transition through miR156-Targeted SQUAMOSA PROMOTER BINDING-LIKE Transcription Factors. Plant Cell 24, 3320–3332.

Zhang, H.Y., Hu, J.H., Qian, Q., Chen, H., Jin, J., and Ding, Y. (2016a). Small RNA Profiles of the Rice PTGMS Line Wuxiang S Reveal miRNAs Involved in Fertility Transition. Frontiers In Plant Science 7.

Zhang, S.X., Liu, Y.H., and Yu, B. (2015). New insights into pri-miRNA processing and accumulation in plants. Wiley Interdisciplinary Reviews-Rna 6, 533–545.

Zhang, W., Xie, Y., Xu, L., Wang, Y., Zhu, X.W., Wang, R.H., Zhang, Y., Muleke, E.M., and Liu, L.W. (2016b). Identification of microRNAs and Their Target Genes Explores miRNA-Mediated Regulatory Network of Cytoplasmic Male Sterility Occurrence during Anther Development in Radish (Raphanus sativus L.). Frontiers In Plant Science 7.

Zhang, Y.-C., Yu, Y., Wang, C.-Y., Li, Z.-Y., Liu, Q., Xu, J., Liao, J.-Y., Wang, X.-J., Qu, L.-H., Chen, F., Xin, P., Yan, C., Chu, J., Li, H.-Q., and Chen, Y.-Q. (2013). Overexpression of microRNA OsmiR397 improves rice yield by increasing grain size and promoting panicle branching. Nature Biotechnology 31, 848–852.

Zheng, C.F., Ye, M.X., Sang, M.M., and Wu, R.L. (2019). A Regulatory Network for miR156-SPL Module in Arabidopsis thaliana. International Journal Of Molecular Sciences 20.

Zhou, X.N., Zhang, Z.H., and Liang, X.H. (2020). Regulatory Network Analysis to Reveal Important miRNAs and Genes in Non-Small Cell Lung Cancer. Cell Journal 21, 459–466.

Zhu, Q.H., and Helliwell, C.A. (2011). Regulation of flowering time and floral patterning by miR172. Journal Of Experimental Botany 62, 487–495.

