## Supplementary figures and images for "MiRNAs profiling and degradome sequencing between the CMS-line N816S and its maintainer line Ning5m during anther development in pepper (*Capsicum annuum* L.)"

### Figure S1

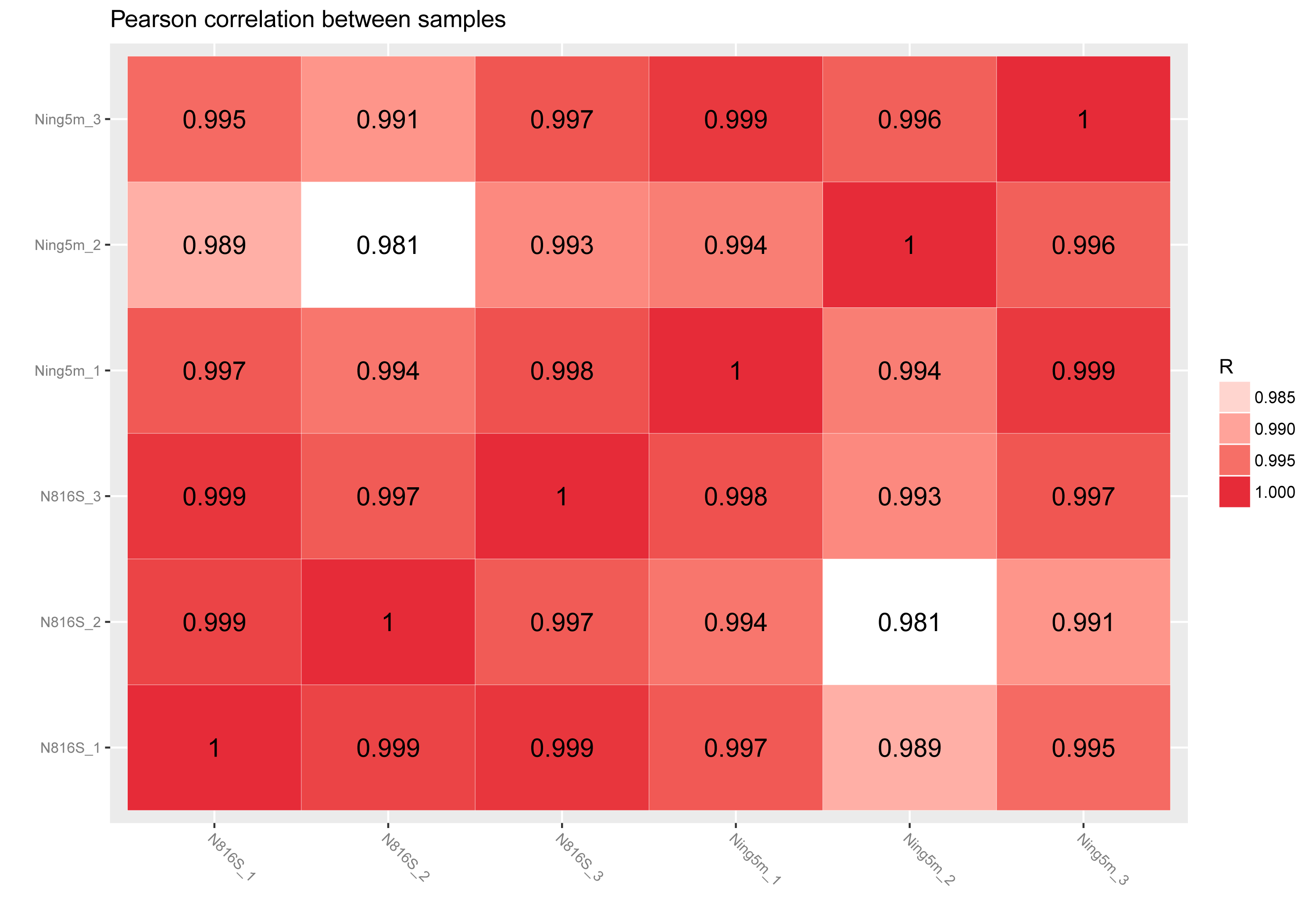
